# Partial coherence and frustration in self-organizing spherical grids

**DOI:** 10.1101/560318

**Authors:** Federico Stella, Eugenio Urdapilleta, Yifan Luo, Alessandro Treves

**Author notes:** Correspondence: Alessandro Treves, Cognitive Neuroscience, SISSA, via Bonomea 265, 34136 Trieste, Italy.

## Abstract

Nearby grid cells have been observed to express a remarkable degree of long-range order, which is often idealized as extending potentially to infinity. Yet their strict periodic firing and ensemble coherence are theoretically possible only in flat environments, much unlike the burrows which rodents usually live in. Are the symmetrical, coherent grid maps inferred in the lab relevant to chart their way in their natural habitat?

We consider spheres as simple models of curved environments and, waiting for the appropriate experiments to be performed, we use our adaptation model to predict what grid maps would emerge in a network with the same type of recurrent connections, which on the plane produce coherence among the units. We find that on the sphere such connections distort the maps that single grid units would express on their own, and aggregate them into clusters. When remapping to a different spherical environment, units in each cluster maintain only partial coherence, similar to what is observed in disordered materials, such as spin glasses.

## 1. INTRODUCTION

What are the defining properties of grid cells? In the 15 years since their discovery in the medial entorhinal cortex (mEC) (Fyhn et al., 2004), two organizational principles appear to have emerged as the cornerstones of the phenomenon. The first one is, of course, the positioning of the fields of each individual cell at the vertices of a regular hexagonal tessellation of the environment (Hafting et al., 2005). The second, a strong propensity of local ensembles of these cells to maintain their co-activation patterns across conditions and environments (Fyhn et al., 2007); in striking contrast to the behavior expressed by neighboring place cells, which make the swapping of activation partners (“remapping”) one of their defining features (Bostock et al., 1991). These two properties, the former expressed at the single-cell level, the latter constraining collective states of activity, have come to be regarded as quintessential to grid cells. They are often yoked together when discussing the “grid cell code” (Yoon et al., 2013; Burak, 2014; Stemmler et al., 2015), thus leaving it unclear whether such code is expressed more in the regularity of field arrangements or in the constancy of spatial phase relations, or in a necessary combination of both.

It should not be forgotten, however, that grid cells have been first described and mostly studied in flat, empty, bounded environments. Their entanglement could be possibly related to the Euclidean geometry of this very specific sort of environment, rather than being intrinsic to the cells. The question, then, is to what extent would single-cell and population properties still co-occur, when such a specific setting is abandoned, to reach for more naturalistic settings of complex, curved, partially open environments, such as the burrows where rats live in the wild (Calhoun, 1962).

In two previous modelling studies, we argued that the notion of the hexagonal grid may need to be generalized in order to predict the behavior of such cells in environments of constant positive or negative curvature. With sufficient exposure to these environments, our model indicates how single grid cells may adapt by producing regular tessellations consistent with the underlying curvature (Stella et al., 2013; Urdapilleta et al., 2015): tessellations with 5-fold or lower symmetry for positive curvature; 7-fold or higher symmetry for negative curvature. An analysis based on Calhoun’s (1962) study leads to the conclusion that the standard 6-fold grid symmetry would arise at the single-cell level only when the curvature is near zero; while the actual range of curvature values of the natural Norway rat habitat extends further, both at the negative and at the positive ends of the spectrum (Fig.1).

**Figure 1.**
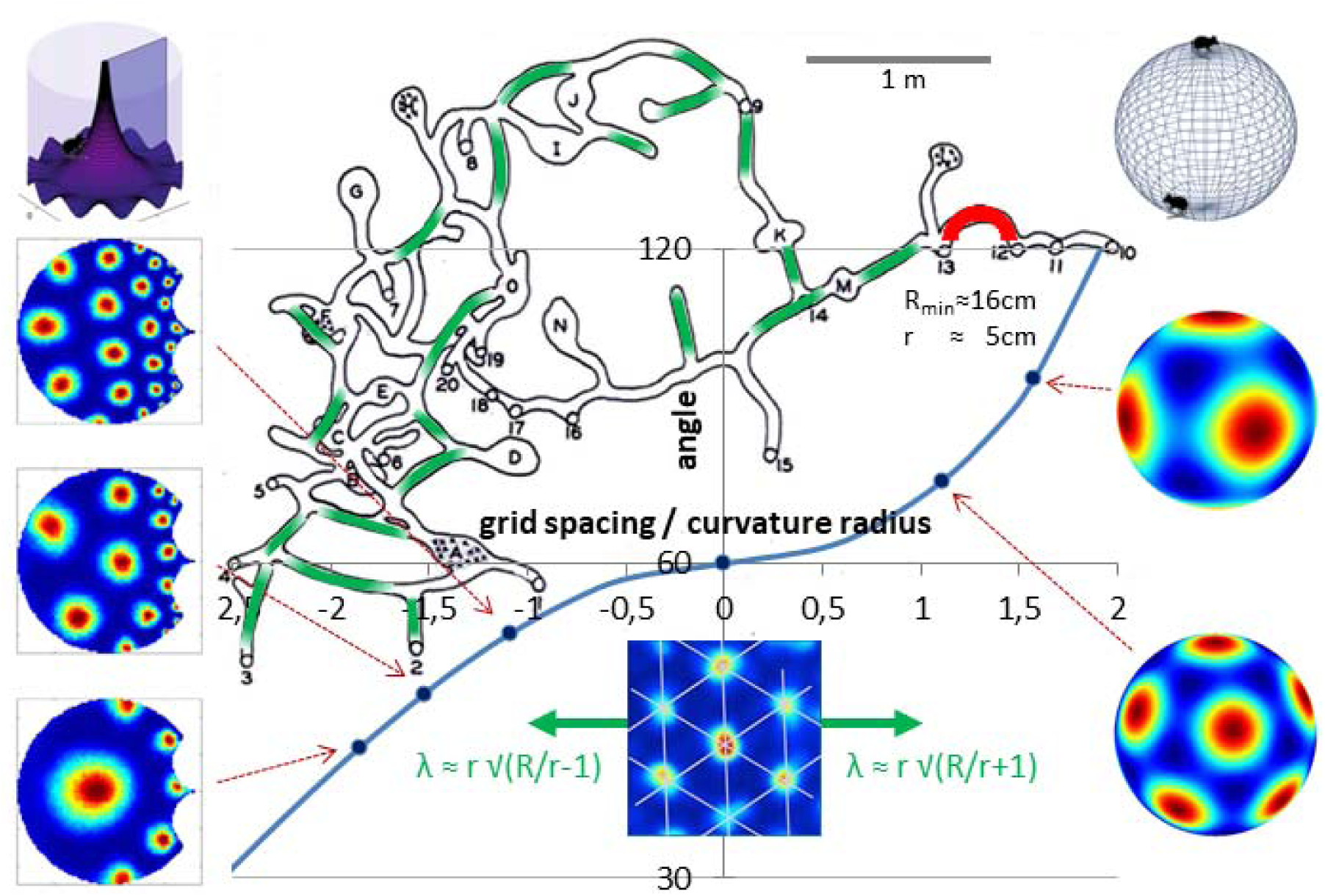
Natural Norway rat environments span limited stretches with negligible curvature. Main graph: the symmetry expected at the single-unit level for different values of constant Gaussian curvature, measured by the ratio between grid spacing s and the radius of curvature λ. Blue curve: theoretical relation between the angle α=2π/n of the n-fold symmetry and the ratio s/λ, cosh(s/λ)=cos(α)/[1-cos(α)] (negative curvature) and cos(s/λ)=cos(α)/[1-cos(α)] (positive curvature). Symmetric arrangements for n=4,5 (on a sphere, right), 6 (on the plane, center) and 7,8,9 (on a pseudo-sphere, left) are indicated, with possible curved environments to be used in the laboratory at the top of the left and right column. Green arrows from the flat 6-fold grid example indicate the range where it may be relevant, |s/λ| ≤ 1, before other symmetries prevail. Superimposed on the graph is a drawing of a Norway rat den, from Calhoun (1962). He estimated the inner radius of the tunnels, r, to be below 5cm, which implies that for the radius of Gaussian curvature λ of a curved tunnel to be of order the grid spacing s, even for a small s=40cm, the outer radius R of the den has to be several meters. This means that the 6-fold symmetry is relevant only to roughly straight tunnel segments, approximately indicated in green, while most of the den (example tunnel in red) does not admit symmetric grids. The chambers are too small to reveal spherical arrangements, and their representation may be more akin to that of the turning points in the hairpin maze (Derdikman et al, 2009).

What about the effect of interactions between grid cells? What was shown in (Stella et al., 2013) is only how a population of non-interacting grid-like units can self-organize a representation of the spherical surface, where each unit ends up displaying an independently “oriented”, often quasi-regular grid. There, grid patterns emerged due to the progressive, unsupervised sculpting of the feed-forward connections through Hebbian plasticity induced by navigation-related activity. Contrary to continuous attractor models (Burak and Fiete, 2009), interactions between mEC units were not needed for the emergence of individual grid maps – possibly, only for their coordination at the population level. Indeed, studies of the same model on *planar* environments have made clear how the introduction of lateral connectivity in the mEC population can induce an alignment among units, resulting in a common orientation of the fields emerging from the feed-forward self-organization process, while also reinforcing their symmetric arrangement (Si et al., 2012; Si and Treves, 2013). Urdapilleta et al. (2015) have observed that also in environments with constant negative curvature the lateral interactions tend to favor a coherent arrangement of the fields across units, but such environments are dominated by their boundaries, which leads to arbitrary modeling choices, that in turn prevent reaching firm conclusions. A complete sphere, on the other hand, has no boundaries, and thus offers a conveniently simple model of a curved environment. Moreover, it has been effective as an experimental set-up, both on the outside (Harvey et al., 2009) and on the inside (Kruge et al., 2013; Kruge, 2016), giving hope that once complemented with the appropriate sensory surround, it will allow developmental studies to investigate the formation of spherical representations in rodents. In the meantime, here we use the adaptation model to study the effect of collateral interactions among grid-like units self-organizing on a spherical world. We also briefly comment on the additional effects expected of gravity, and of boundaries, when extending the analysis from our artificial spherical environment to ecologically plausible ones – an extension that we leave for future studies.

## 2. THE MODEL

The basic details of the model are identical to (Stella et al., 2013), and are described in Appendix 1, with the critical addition of a set of recurrent collaterals connecting units of the EC layer.

Time is discretized in steps of length *t* = 0.01s. The total length of a simulation is of 100 million steps (corresponding to nearly 12 days of continuous running, a very long time, to ensure that the self-organization process has approached its asymptote). The virtual rat moves on the surface of a sphere of radius 52.6cm with a constant speed of *v* = 40cm/s. To obtain *smooth* random trajectories, resembling those observed in experiments, running direction changes gradually after each step, resulting in an extended correlation over time. For simplicity, the change in running direction between two consecutive steps of the virtual rat is sampled from a Gaussian distribution with zero mean and standard deviation *h* = 0.2 radians. The virtual rat always runs along the great circle determined by its running direction. Our model is comprised of two layers. The input array represents e.g. the CA1 region of the hippocampus and includes *N*_hipp_=1400 units with their fields regularly arranged to evenly tile the spherical surface. The output network is comprised of a population of *N*_mEC_ = 250 would-be grid units, all with the same adaptation parameters – hence they represent a single mEC module, in relation to the modules discovered by Stensola et al. (2012).

In a limited set of simulation intended to explore the effects of boundaries, and of gravity, we used hemispheres instead of full spheres. The effects of a boundary, corresponding to the equator, was assessed both with isolated hemispheres, in which case the boundary was reflecting the trajectory of the virtual rat, and with hemispheres embedded in a flat surround, where the boundary amounted only to a sudden change of intrinsic curvature. Without gravity, whether the hemisphere is concave or convex makes no difference. We also simulated trajectories on concave and convex hemispheres with gravity, and the latter was modelled by an additional force dragging the trajectory towards the equator (in the convex hemisphere) or the bottom of the bowl (in the concave one). This force was parametrized by the change in speed when moving downward and the strength of the downward pull applied to the vector expressing the current direction of motion (see Appendix 1).

Similarly to the planar case, we introduce collateral weights between mEC units to induce the coordinated development of their firing maps. The weights are set in the following way: each unit is temporarily assigned a preferred position, an “auxiliary field” at coordinates (ϕ,θ) on the sphere, and a “preferred direction” (angle relative to the meridian, with 0 pointing towards the North Pole). The coordinates as well as the angle are randomly chosen. These auxiliary fields are introduced solely to define the collateral weights, and not to position the subsequently developing grid fields, nor do they play any role in other parts of the simulations. They are only used, in other words, to induce a notion of similarity among output units. The collateral weight between unit *i* and unit *k* is then calculated as

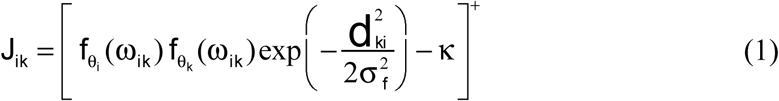

where []+ denotes the Heaviside step function, κ=0.05 is an inhibition factor to favor sparse weights, *f* is a tuning function described in the Appendix 1 and ω_*ik*_ is the direction, with respect to the North Pole, of the line connecting the auxiliary fields of unit *i* and *k*, along the great circle. σ _*f*_ = 10cm denotes how broad the connectivity is, and *d*_*ki*_ is defined as

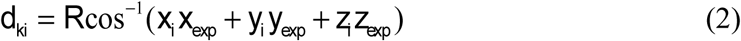

i.e., it is the distance between the coordinates of the auxiliary field (*x*_*i*_, *y*_*i*_, *z*_*i*_) and the *expected* position of a virtual rat that had started at the auxiliary field of unit *k* and had moved 10 cm along the geodesics joining both fields, corresponding to 250 ms of reverberatory delayed activity of movement along this direction. The definition of the weights leads to a localized connectivity pattern, such that strong positive interactions are only generated between units with similar preferred head direction and activation fields appropriately shifted along the same head direction (Kropff and Treves, 2008; Si et al., 2012). The resulting connectivity is rather sparse, with only about 8% of the possible pairs sharing a non-zero weight. As with the feed-forward connectivity, normalization on this set of connections is performed by setting a unitary **L**^2^ norm on the pre-synaptic strengths for each mEC unit. Moreover, the relative strength of the recurrent input with respect to the feedforward input was reduced to 0.2 for most of the simulations (see Appendix 1). Notice that our model does not include plasticity on the recurrent set of connections: their value is defined once, at the beginning of each simulation, and then kept unmodified throughout. Note also that with a radius R=52.6cm and the adaptation parameters we use, most units tend to have 12 fields in the simulations with recurrent connections, but 13 or 14 fields without (see Fig.S1, top). Still, we choose to use the same value of R in the two cases for ease of comparison. When separately varying R in order to optimize the proportion of units evolving 12 fields in each condition, and taking into account all units, the simulation without collaterals yields maps much closer to the “soccer ball” ideal (Fig.S1, bottom).

## 3. RESULTS

Simulations produce units with different number of fields, as shown in Fig.S1. In the following, we set the radius to the same value R=52.6cm and consider only units which have developed 12 fields. We compare simulations with and without lateral connections. For each simulation, the same set of grids are left to self-organize in two distinct, independent environments.

### Grid Map Distortion

At the single-map level, the introduction of lateral connections has the immediate result of interfering with the development of a regular grid structure. We can quantify this phenomenon by computing the spherical correlation (as the best match over any possible rotation) of the activity map developed by each unit with the template of a perfect 12-field “soccer ball” rate map (Stella et al., 2013). One sees a marked effect of recurrent interactions as a general increase in the distance from the best-matching template, even though the radius of the sphere can be adjusted so that even with the recurrent collaterals most units produce 12 fields (Fig.2C,D, and Fig.S1). The simulations without collaterals produce fairly good exemplars of an ideal spherical grid. This is not the case when recurrent collaterals are introduced: the interactions among EC units lead to a disruption of the regular arrangement of their fields (see the examples in Fig.2A,B; and the quantitative measures in Fig.S2).

**Figure 2.**
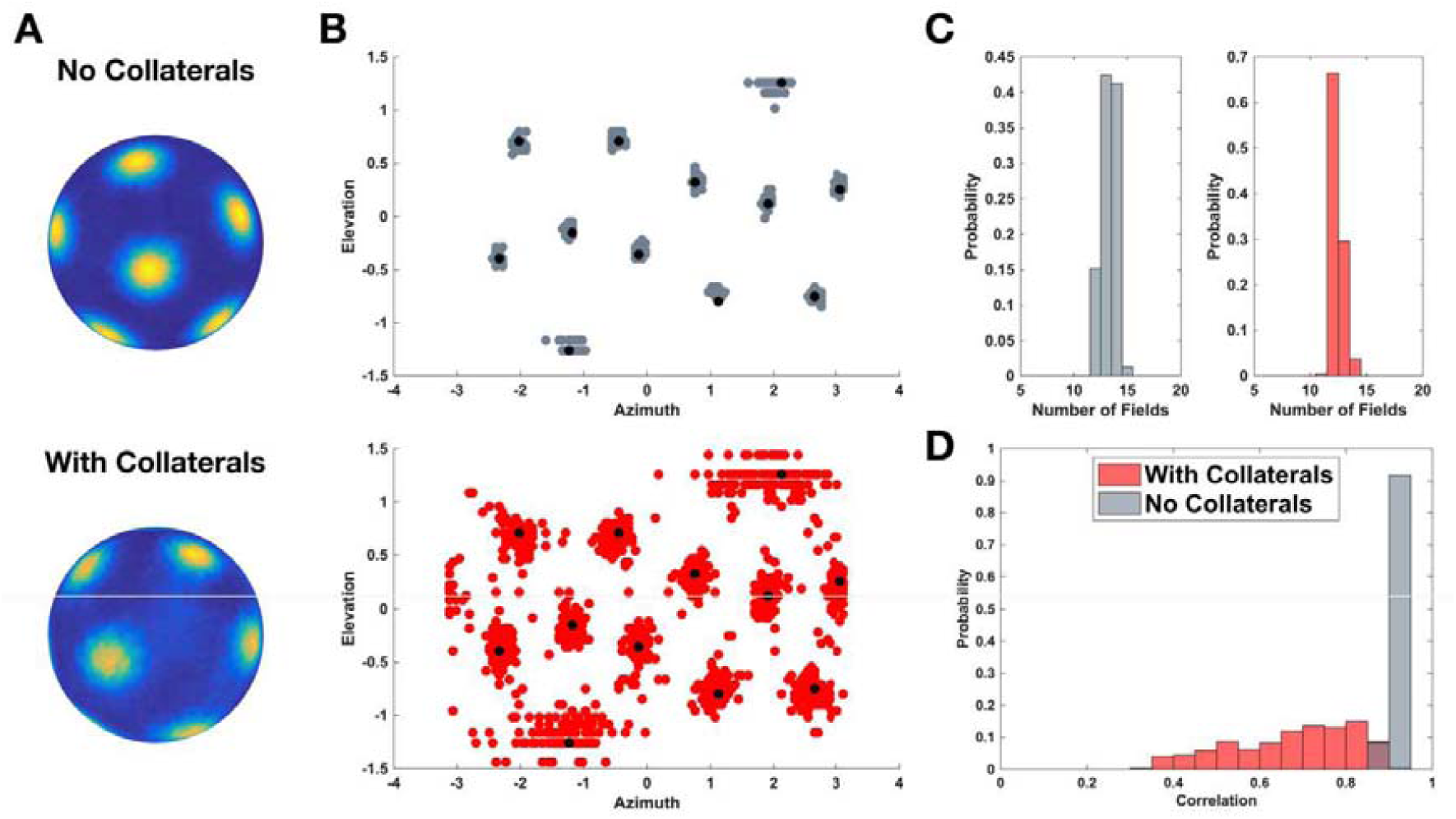
Collateral interaction distorts the grid pattern on a sphere. **A** Two representative examples of activity developed by grid units on the sphere. Top: unit from a simulation without collateral connectivity. Bottom: unit from a simulation in which units interact through collaterals. **B** Distribution of the position of all the fields from a population after the rate map of each unit has been rotated to maximize its overlap with a common “soccer ball” template (the field centers of this perfect grid are shown in black). Top: fields of a population of non-interacting units (mean distance from perfect center: 3.52°); Bottom: fields of a population of interacting units (mean distance from perfect center: 7.96°). **C** Distribution across the population of the number of fields developed by units, with R=52.6cm. Left: no collaterals; Right: with collaterals. Sample sessions. Values across sessions: mean fraction with 12 fields, for no interaction 0.13, with interactions 0.68. **D** Correlation of all units with 12 fields with a best-rotated ‘soccer ball’. Aggregate from all sessions.

This illustrates how quickly the spherical case departs from what is observed on a planar surface. There, it has been shown (Si et al., 2012; Si and Treves, 2013), the presence of lateral connections has the crucial role of inducing a common orientation in the grid population and does not hinder, in fact enhances, the quality of the grids developed by the system. The same process does not occur here, where a similar attempt to induce coordination in grid-evolving units appears at odds with the regularity of the grids.

### Grid units tend to cluster

One can then ask, what are the effects of connections on the whole ensemble of grid units, and what does the “common orientation” that they should induce look like, on the sphere? We answer these questions by analyzing the spatial (spherical) similarity in the structure of the maps developed by different units. To do so, for each unit, we computed the activity auto-correlation after 373,248 different rotations. Rotations were randomly drawn to evenly span the space of possible Euler rotations (2π×π× 2π, considering also the cosine factor). We then sorted the rotations according to their autocorrelation score, from highest to lowest. We used the first 5000 best rotations to compute their distribution density in the 3-dimensional space of Euler rotations Γ_*i*_ (ϕ,θ, Ψ) (where *i* denotes the unit). For each pair of units (*i,j*) we then computed the overlap between Γ_*i*_ (ϕ,θ, Ψ) and Γ_*j*_ (ϕ,θ, Ψ) (as a correlation). This correlation (or effectively, distance) matrix was used to identify clusters of units sharing a similar rotation distribution. The clustering was performed with the “ward” algorithm and the number of clusters was optimized over the range [4-15].

The outcome of this clustering algorithm shows how the population effectively breaks down into sub-groups of segregated units, each developing an internal degree of coherence that is higher than that shared by the entire population. In Fig.3, the spatial position (on a 2D projection of the sphere) of the field centers of all units with 12 fields in the population are shown colored according to their cluster membership. To a large extent, each of these clusters expresses a common orientation, meaning that each of the 12 fields of a unit tends to appear grouped with those of every other unit in the cluster. Different clusters appear instead unrelated, roughly to the same extent that individual units are in the non-connected simulations (Fig.3B).

**Figure 3.**
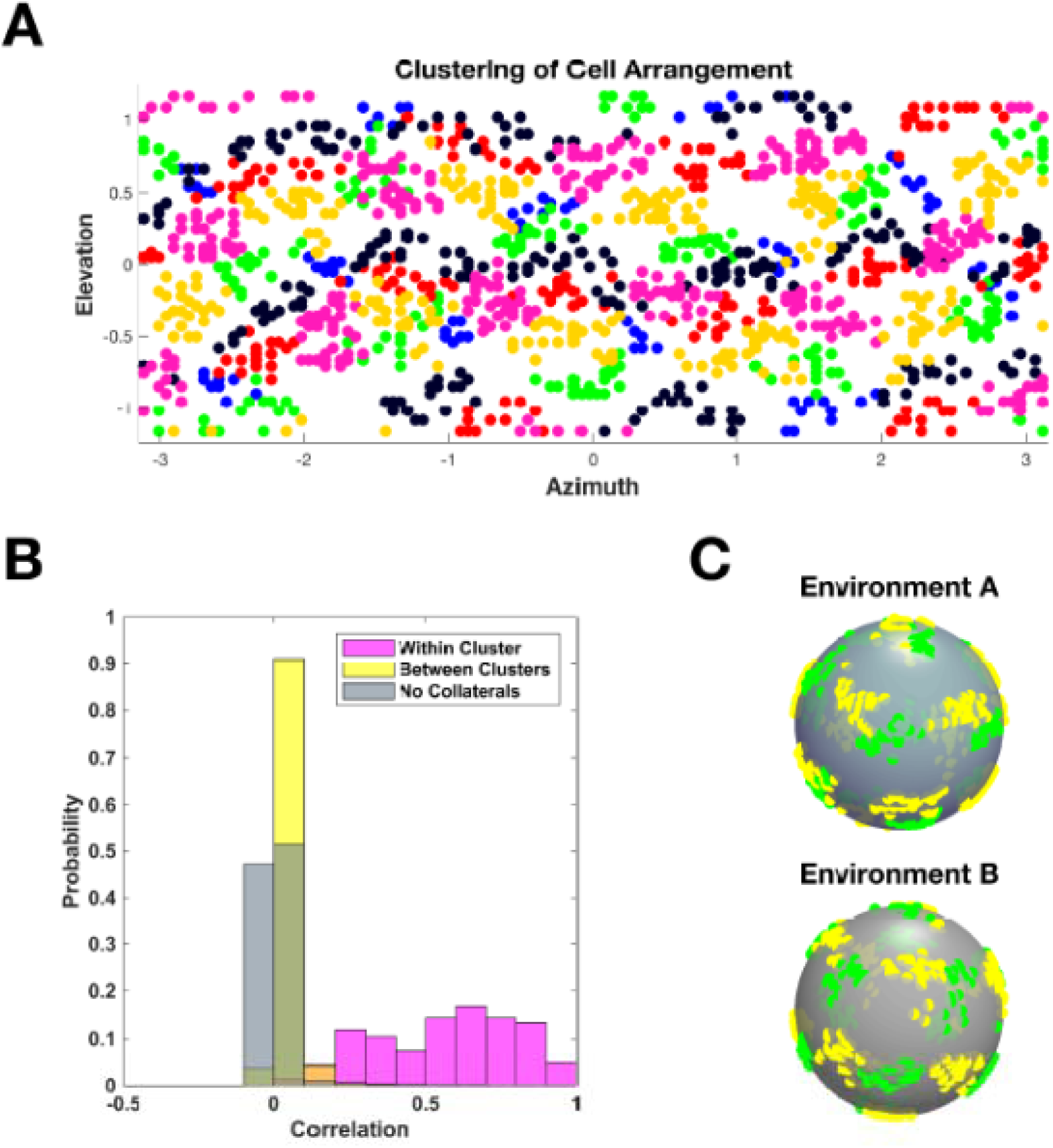
Interacting grid units tend to cluster. **A** Spatial distribution of the fields of a population of grid units over the surface of a sphere. Each field center of each unit is assigned a color according to which cluster the unit belongs to. Sample session. Average number of clusters across sessions: 7.6. **B** Measure of clustering quality: distribution of pairwise unit correlations between the 3D density of their 5000 best rotations in the space of Euler angles (the measure used to define clusters). Sample session. **C** Example of the arrangement of the fields from the units of two clusters (green and yellow, as in A). The positions of their fields are shown in two different spherical environments. Clusters were defined solely in the first environment.

Crucially, the membership to one cluster is a feature that is carried over to new environments almost entirely unaltered (we describe the remapping procedure in the next paragraph). In Fig.3C we show a similar plot of the field centers, this time for only two of the clusters, to highlight their correspondence in two different environments. The grouping is conserved, as is the mutual avoidance of the fields in the two clusters. Thus we observe how on the sphere, the interaction between units results into a break-down of the population, with different sub-sets acquiring a coordinated arrangement, while at the same time it forces each unit to distort its firing pattern away from that of an ideal grid. These features are consistently reproduced across environments.

### Remapping

In order to gain a better understanding of the mechanisms underlying grid formation and of the spatial code they can generate, we need to address the properties of remapping. To that end, maps were developed independently in two environments: the set of place cell inputs and the associated feed-forward connectivity was randomly initialized in each of the two environments. Only the strength of the recurrent collaterals (when present), and thus of the grid cell interactions, was maintained after remapping. For each unit, we compare the maps developed in the two environments, maps A and B. Taking map A, we again apply a large random set of rotations spanning the entire Euler rotations range, and for each rotated version of map A we compute the resulting overlap (correlation) with map B. We first consider only the rotation associated with the highest similarity score – the “best” rotation – and its associated rotated version of the map, A’.

Results are shown in Fig.4A: in red one can see the distribution of correlation values between B and A’, and for comparison the results for the best rotation of a perfect grid (in light brown). The two distributions lie in the same range of high correlation, although the perfect grid can usually be rotated to achieve a higher correlation with the map in B, suggesting that the distortion observed in A is independent of that in B. We can compare both distributions with that obtained by correlating for each unit the best rotated map A’ with the map of each other unit in the same cluster, in B (magenta). In this case correlation values are somewhat lower but, consistent with the partially coherent behavior expressed by units in the same clusters, they are still significantly higher than those obtained using either the maps in B of units in other clusters (yellow) or of all the other units (not shown).

Next, we observe that, if the two maps were perfect soccer balls, there would be 12*5=60 equivalent ways to rotate one into the other. Because of the distortions from the most symmetric configuration, the degeneracy is only approximate, but still massive: there are many different rotations, spanning a diverse set of Euler angle triplets, that lead to almost the same correlation values as the best rotation, for each unit. In fact, if we take for each unit its N_Best_ rotations of the map in A, we see that their average correlation with the corresponding map in B is a smooth function of N_Best_, which averaged over the population shows a significant decrease only when N_Best_ reaches into the thousands (among an arbitrarily set range of 373,248 randomly chosen triplets (ϕ,θ, Ψ) of Euler angles; Fig.4B).

**Figure 4.**
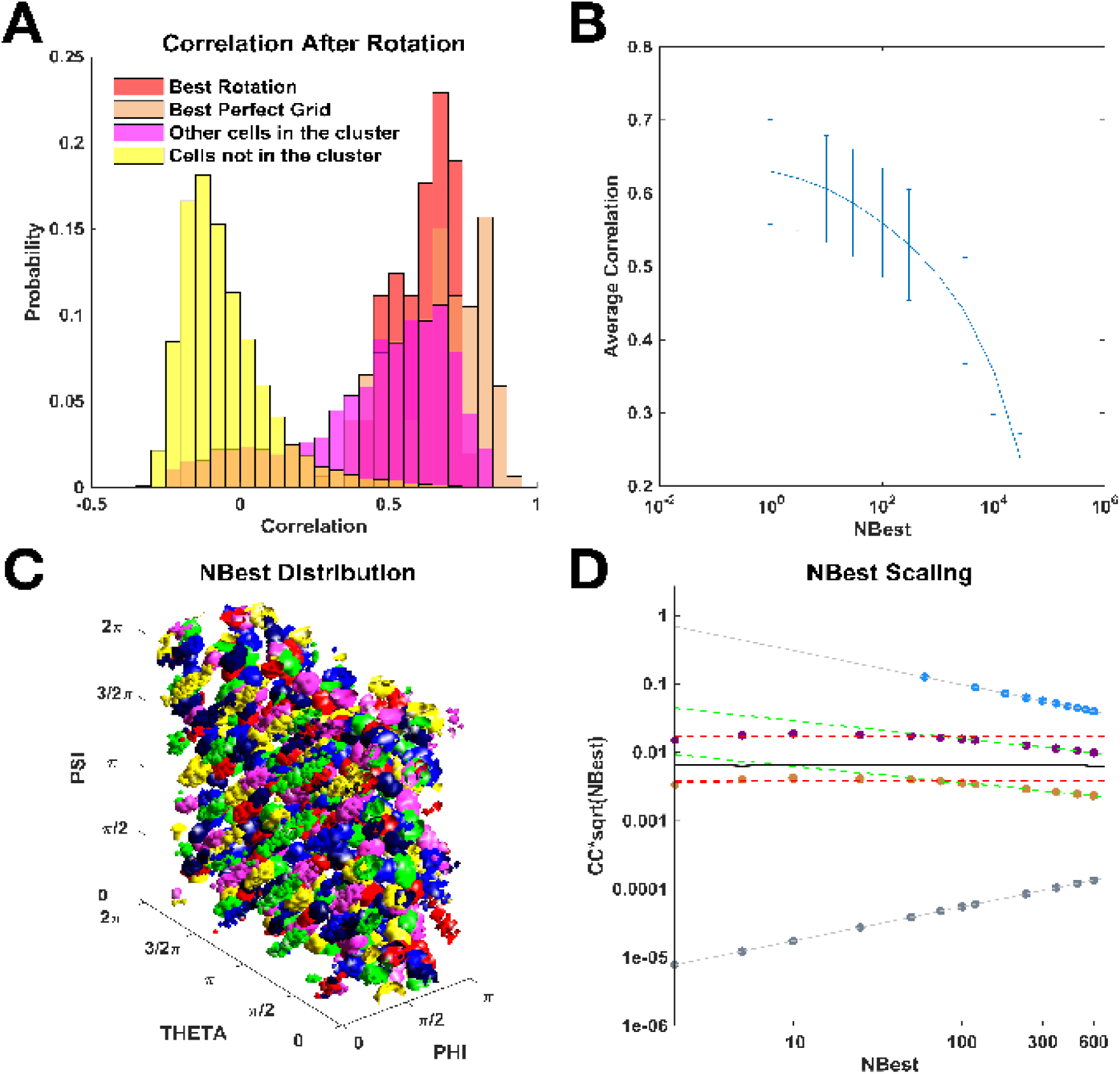
Remapping preserves the clusters. **A** Distribution of overlaps after the best rotation (out of 373,248 randomly drawn rotations). All sessions. Mean overlap: with the map in B of the same unit, 0.615; with those of other units in the cluster, 0.455; with those of units not in the same cluster, -0.050; with those of all units, 0.006 (not shown). Mean overlap of the best rotated perfect grid with the maps in B, 0.659. **B** Correlation-N_Best_ dependence. Average value of the correlation between the map in B and the map of the same unit in A rotated N_Best_ times (bars denote SDs). **C** Example of the spatial density distribution of best rotations for cells belonging to different clusters. Each color represents the distribution of the N_Best_=500 angles for 5 random units for each cluster. Sample session. **D** N_Best_ scaling and partial coherence. Different scaling CC=1/N_Best_^β^ of the Clustering Coefficient (computed over distributions like the one in C) for different types of remapping. Logarithmic scales. For clarity, the quantity on the y-axis is CC·N_Best_^1/2^. Grey: random remapping of non-interacting units, β=0. Blue: coherent remapping, β=1. Purple: CC computed using interacting grid units from the same cluster. Brown: CC computed using all units in an interacting population. Dashed black: β=0.5; dashed red: CC computed using N_Best_ up to 100 (average over simulations: β=0.53 all units, β=0.52 units in same cluster); dashed green: CC computed using 100 < N_Best_ <= 700 (average over simulations: β=0.74 all units, β=0.71 units in same cluster); Black: computed using numerical estimation of the analytical formulation (see Appendix 2).

Remarkably, the clusters of units defined on sphere A maintain a *partial* coherence once remapped onto sphere B, as already suggested by the example in Fig.3C. If we randomly choose 5 sample units per cluster, and consider their N_Best_=500 individually most correlated rotations, we find that they cluster into distinct ‘islands’ in Euler space, with each cluster contributing 60 regularly arranged islands, as shown in Fig 4C. Inside each island, however, chaos prevails. Some single units contribute many more of their best rotations to particular islands, and the shape of each island appears randomly distorted.

After failing to observe any further geometrical structure within the islands, we resorted to a quantitative measure of the extent to which the best rotations are concentrated across units. We define a clustering coefficient, CC (Cerasti and Treves, 2013), that measures, starting from the 373,248 randomly chosen Euler triplets and taking the N_Best_ distinct rotations for each of N units, the probability that two such triplets coincide (see Appendix 1 for the definition). For a perfectly coherent rotation the N_Best_ rotations would be the very same triplets across units, hence CC=1/N_Best_. For a totally incoherent remapping, triplets would coincide at chance level, hence CC=1/373,248. Fig.4D shows that, whether we take only units within the same cluster or in the entire population, the clustering coefficient has intermediate values, scaling approximately as 1/√N_Best_, for N_Best_ small – corresponding to a horizontal line in Fig.4D. For larger N_Best_, a steeper decrease prevails, presumably because the population remains less than fully coherent also when allowing for looser remapping correspondence. In Appendix 2 we show that an intermediate scaling can be expected from a simple analytical model. A direct numerical evaluation of the mean field formulation for the clustering coefficient (see Equation A2.4 in the Appendix 2) yields a behavior in very good agreement with the one observed in simulations (Fig 4D, black continuous line). Here the deflection from the 1/√N_Best_ scaling appears to happen for larger values of N_Best_ (lying outside the right margin of the plot), presumably as an effect of the mean field approximation. The model does not seem to predict, however, an exact square root scaling, and it remains unclear to us whether the exponent β≈1/2 that we find to characterize CC≈1/N_Best_^β^ (for N_Best_ small) is fundamental or a mere coincidence.

An extensive set of simulations with varying overall interaction strength among the units (through a prefactor) indicates that the absolute value of the clustering coefficient depends mildly on the prefactor, but its scaling exponent β≈1/2 for N_Best_ small is the same (not shown), and the clustering coefficient is constant at 1/373,248 only when the prefactor is strictly zero.

### Towards ecological plausibility

The full spherical environment may be approached with an appropriate experimental set-up (Mayank Mehta, personal communication; by using spherical virtual reality, see also Aghajan et al., 2015), but is far from those in which rodents have evolved in the wild. The sphere does not include several features of e.g. the systems of burrows rodents dig as their homes. To begin considering the relevance of such features to grid maps, we start here with two: the presence of boundaries and the pull of gravity. By altering exploration and navigation behavior, both these features are expected to have at least an indirect influence on spatial codes, also in curved environments.

The effect of a *boundary* can be appreciated already by simply slicing a sphere in two halves, and running simulations of non-interacting units in one hemisphere (Fig.S4A,B). If the cut were to be randomly oriented with respect to a perfect soccer-ball grid, one would expect definite proportions of the two hemisphere patterns in Fig.S4A. In particular, the bottom arrangement with 3 fields around the pole should occur with *q*=28.6% of the units, as can be calculated from the exact formula

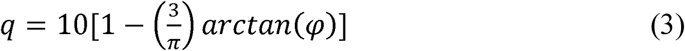

here φ=1.618.. is the golden ratio. In simulations, however, it occurs with about half that frequency, *q*=14±5% (Fig.S4B), as the fields of individual units tend to form away from the border. This also distorts the “pentagonal gridness” of each map on the hemisphere. Further, the presence of a hemispheric bump or cavity in an otherwise flat environment distort the hexagonal gridness of the fields near the boundary with the hemisphere. We have quantified this effect by running simulations on a flat ring which may contain either a circular hole or a hemisphere, and measuring the standard grid score on the flat part of would-be grid units, this time interacting through recurrent collaterals. The presence of the (curved) hemisphere halved the average score, with respect to simulations run around a hole, from 0.39±0.16 to 0.19±0.07 (Fig.S4C,D).

Further grid distortions appear if gravity is present. In the adaptation model, they are due to the unequal exploration of different latitudes on the hemisphere, which obviously deviates in opposite directions from an even sampling, depending on whether the hemisphere is set as a hill or as a valley. In Fig.S5 we show examples of simulations run on a hill, in which trajectories were determined by adding to the standard algorithm, generating a random movement vector at each time step, a downward bias in speed (40% faster) and in turn selection, to model the downward gravity pull (see Appendix 1 for details). As can be seen, grid fields tend to cluster around the equator (Fig.S5A,B). Interestingly, the resulting decrease in correlation with the perfect soccer ball field distribution is similar for non-interacting grids (Fig.S5C) and for interacting ones (Fig.S5D), which as discussed above already deviate more from the perfect arrangement also in the absence of gravity.

## 4. DISCUSSION

The simulation of our adaptation model, allowing for collateral interactions among the units, indicates a radically different nature of the grid code on the sphere. The same interactions which on a plane suppress fluctuations and lead to a collapse into a smooth continuous attractor recruiting all units with the same or similar grid spacing (Si et al., 2012; Urdapilleta et al., 2017), on a sphere lead to a hierarchy of effects on single-unit maps that

a. are distorted from the available symmetric ‘soccer ball’ field arrangement
b. are forced into clusters of units with approximately overlapping fields
c. remap to a different sphere with only partial coherence, even within clusters.

We argue that such effects are due to the unresolved conflict between regular tessellation at the single unit level (due to adaptation dynamics) and global coherence at the population level (induced by recurrent connections). Unlike the planar case, where these two aspects can coexist in the same grid code, spherical geometry only allows for a compromise solution, where both regular tessellation and population coherence are only partially attained. We regard these as predictions that could be validated or falsified by experiments which are doable in rodents, even though they may require *ad hoc* arrangements to allow for the slow emergence, possibly only during a two-week developmental period, in rats (Langston et al., 2010), of a stable set of 2D maps.

Alternative models of grid map formation may lead to different predictions, but we would not know how, and are not currently aware of attempts by others, to extend existing models e.g. the oscillator interference model (Burgess et al., 2007) or the continuous attractor model (Burak and Fiete, 2009), to work on a sphere. Crucially, the continuous attractor model is based on the compatibility, contingent to Euclidean (in 2D, planar) spaces, of a single-unit hexagonal pattern, potentially extended to tile environments of any size, with congruent phase-offsets in different units. While this model can account for several grid cell properties and for their rapid manifestation, as observed in laboratory experiments, it appears that its theoretical premises make it inapplicable to environments of non-zero curvature. The regular phase off-set that allows to project the activity from the ‘cell layer’ envisaged by the model onto real space is just not possible with spherical geometry, leading to a loss of coherence in the activity of different units. It is also unclear to us how the oscillator interference model could be applied to curved environments.

A sphere is of course an even more artificial rearing environment than a flat box, but we believe that it may help capture a fundamental trait of grid cell coding, by pointing at those properties of grid cells often assumed to be universal but in fact stemming from the use of flat, bounded environments. The qualitative characteristics a)-b)-c) may be general to any curved environment, and they can be contrasted with the character of grid cell activity in rodents reared in standard laboratory conditions. In this sense, a sphere may be closer than a plane to the ecological condition of a Norway rat system of burrows (Calhoun, 1962). Grid cell representations may be presumed to have evolved to be relevant to rodents living in the wild.

In humans, the same fMRI hexagonal signature that has been hypothesized to reflect grid cell activity in a virtual reality navigation task (Doeller et al., 2010) has later been reported when subjects ‘move’ in a 2D space of drawings (Constantinescu et al., 2016), raising the issue of whether hexagonal symmetry may characterize even abstract conceptual spaces, when described by assigning two dimensions (Bellmund at al., 2018). Our model suggests that this may occur only around locations that are either flat *a priori*, or where curvature has been ironed out, perhaps by extensive training.

From a complex systems point of view, it is remarkable how curvature opens up a scenario different from that of a strictly regular, periodic 2D tessellation. In a separate study, we have already argued how the extension of such 2D tessellation to a 3D crystal, a scenario potentially relevant to bats and other animals navigating through 3D volumes, is in fact implausible, because of time scales involved (Stella and Treves, 2015). Here, we make the case that also navigation on 2D manifolds embedded in 3D Euclidean space (such as tree-branches or multi-store buildings) might be associated with a ‘broken’ grid cell representation, retaining only part of the planar symmetry.

In the new scenario, a network of grid units ‘behaves’ more like a disordered system than like a crystal. The approximate inverse square root scaling of the clustering coefficient of the rotations, under remapping, reminds us of the partial coherence of a physical system with impurities, where some interactions are perforce ‘frustrated’ (Mezard et al., 1987). When interactions are short-range, local coherence may survive, avoiding the impurities, somewhat like grid maps away from objects placed in a flat environment (Hoydal et al., 2018; Boccara et al. 2019). A prevailing non-zero curvature is more akin, however, to a system with impurities and long-range interactions, where disorder affects even the shortest organizational scale. Such systems, not unlike human society, can offer only partial coherence.

Partial coherence, together with the realization that a network of grid cells may be endowed with a significant storage capacity (Spalla et al., 2019), in a sense brings back grid cells to the fold of memory systems, next to the place cells, with their multiple charts (Samsonovich and McNaughton, 1997; Battaglia and Treves, 1998). For years, it has been thought that grid cells may afford long-distance path integration (Fuhs and Touretzky, 2006; McNaughton et al., 2006). Partial coherence, however, limits accurate path integration to short distances. Together with the emerging observation that mEC outputs to the cortex, mainly from layer Va, include virtually no grid-cell signal (Alexei Egorov and David Rowland, personal communication), this weakens the theory that mEC operates as a sort of spatial computer, and suggests instead that grid maps are one input that helps set up the spatial component of hippocampal memory representations. Alternative sets of coactivity relations stored on the same synaptic connections, as well as curvature, act on the currently active grid representation as ‘quenched’ disorder, and coexisting with such spin-glass-like disorder appears to be the ultimate challenge for memory systems in the brain (Treves, 2009).

## Acknowledgements

We are grateful to Matteo Toso, who modeled the habitat of the Norway rat, to YuQiao Gu, who carried out some of the early simulations, and to colleagues in the EU ‘Gridmap’ and HFSP RGP0057/2016 collaborations, which also partially funded this study. This report was in part drafted during a 2018 visit by AT to the KITP, thereby also supported in part by NSF Grant No. PHY-1748958, NIH Grant No. R25GM067110, and the Gordon and Betty Moore Foundation Grant No. 2919.01.

## Appendix 1: Model

Additional aspects of the model, besides those reported in the main text:

The input to mEC unit *i* at time *t* is given by

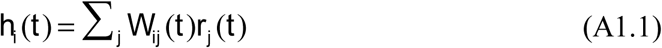

The weight *W*_*ij*_ connects input unit *j* to mEC unit *i*. We assume that at the time the mEC units develop their maps, spatially modulated or place cell-like activity is already present, either in parahippocampal cortex or in the hippocampus. The network model works in the same way with any kind of spatially modulated input, but the place-cell assumption reduces the averaging necessary for learning. Each input unit activity in space is modelled as a Gaussian place field centered at preferred position *x*_*j0*_

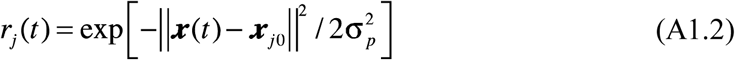

where ***x***(t) is the position at time *t* of the simulated rodent, σ_*p*_ =0.05m is the width of the field and ||***a***-***b***|| is the great-circle distance on the sphere.

### Single Unit Dynamics

The firing rate ψ_*i*_ (*t*) of mEC unit *i* is determined by a non-linear transfer function

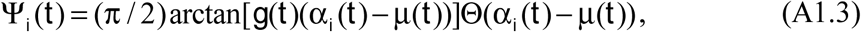

which is normalized to have maximal firing rate equal to 1 (in arbitrary units), while Θ(□) is the Heaviside function. The variable μ(*t*) is a threshold while α*i* (*t*) represents the adaptation-mediated input to unit *i*. It is related to *hi* (*t*) as follows:

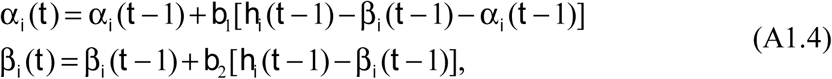

where β*i* has slower dynamics than α*i*, with *b*_2_=*b*_1_/3, *b*_1_= 0.1 (in a continuous formulation, the *b* coefficients become rates, in units of (Δ*t*)^-1^). These adaptive dynamics make it more difficult for a neuron to fire for prolonged periods of time, and correspond to the kernel *K* considered in the analytical treatment (Kropff and Treves, 2008). The gain *g*(*t*) and threshold μ(*t*) are iteratively adjusted at every time step to fix the mean activity *a* = ∑_*i*_ ψ_*i*_ (*t*)/ *N*_*mEc*_ and the sparsity *s* = (∑_*i*_ ψ _*i*_(*t*))^2^ /(*N*_*mEc*_ ∑ ψ (*t*)^2^) within a 10% relative error bound from pre-specified values, *a*_0_=0.1 and *s*_0_=0.3 respectively. If *k* is indexing the iteration process:

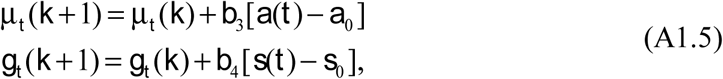

*b*_3_=0.01 and *b*_4_=0.1 are also rates, but in terms of intermediate iteration steps. *a*^*k*^ and *s*^*k*^ are the values of mean activity and sparsity determined by μ_*t*_ (*k*) and *g*_*t*_ (*k*) in the intermediate iteration steps. The iteration stops once the gain and threshold have been brought within the 10% error range, and the activity of mEC units are determined by the final values of the gain and threshold.

### Synaptic Plasticity

The learning process modifies the strength of the feed-forward connections according to a Hebbian rule

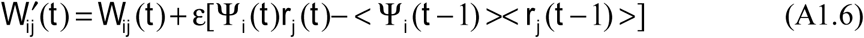

with a rate ε=0.002. <Ψ_*i*_(*t*)> and <*r*_*j*_(*t*)> are estimated mean firing rates of mEC unit and place unit *j* that are adjusted at each time step of the simulation

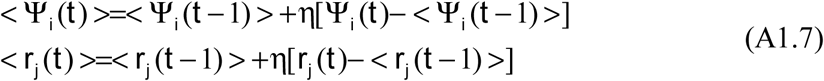

With η=0.05 a time averaging factor. After each learning step, the *W’*_*ij*_(*t*) weights are normalized into unitary norm

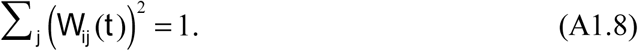

#### HD Input

Head Direction on the sphere is defined as the angle between a vector and the vector pointing towards the north pole. With the addition of HD modulation and collateral connections, the overall input to unit *i* for the interacting case is:

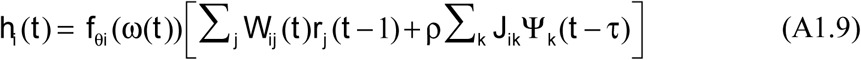

with ρ=0.2 a factor setting the relative strength of feed-forward *W*_*ij*_(*t*) and collateral weights *J*_*ik*_, and τ =25 steps a delay in signal transmission, as discussed by (Si et al. 2012). The multiplicative factor *f*_θ*i*_(ω(*t*)) is a tuning function which is maximal when the current direction of the animal movement ω(*t*) is along the preferred direction θ_*i*_ assigned to unit *i*

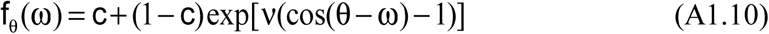

where c=0.2 and ν=0.8 are parameters determining the minimum value and the width of the cell tuning curve. Preferred head directions are randomly assigned to mEC units and they uniformly span the 2π angle.

#### Clustering Coefficient (*CC*)

Given *q* cells and taking the first N_best_ Euler rotations from each, the clustering coefficient was defined as:

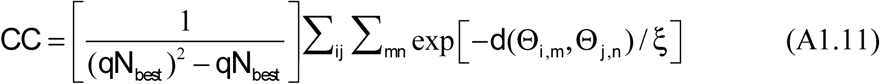

with *i*≠*j* and where *d*(Θ_*i,m*_, Θ _*j,n*_) is the distance between two three-dimensional rotations; and ξ is set to 5 degrees so that only nearly coinciding rotations contribute to the sum.

### Grid field definition and properties

Individual fields for each developing unit were identified as continuous portions of the spherical surface where the unit firing rate was above two times the average firing rate computed over the entire environment. Field size was defined as the number of bins passing the threshold in each continuous region, field height as the maximum firing rate within the continuous region, and the field ellipticity as the ratio between the radii of a circle circumscribed to the field and a circle inscribed in the field.

### Effects of gravity on the animal trajectory

To simulate the change in movement statistics due to the presence of gravity, we modulated the generation of random trajectories in two ways.

1. We assumed that downward movement is executed at greater speed than upward movement. Therefore the animal speed was modulated depending on its running direction as *vg* = *v*(1 – ζ cos(θ_*z*_)) where θ_*z*_ is the angle between the direction of motion and the vertical axis (0 when pointing upward), and ζ is a parameter regulating the strength of the gravitational effect.
2. We also applied a constant, downward pull to the direction of motion by applying a bias in the step-wise choice of a new running direction. Down-ward turns were favored by implementing a Metropolis Markov Chain that only accepted up-ward turns with a certain probability. Namely, a turn was rejected when *u* < α where *u* is a uniform random number on [0,1] and 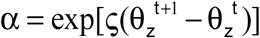. θ_*z*_ is the angle with respect to the z-axis (0 when pointing completely up-ward), ζ is a parameter regulating the strength of the bias, taken here to numerically coincide with that of point 1).

## Appendix 2: Statistical Analysis

Let us consider N units that have developed grid representations on a sphere A, and now develop also grid representations on another sphere B. For every triplet of Euler angles (ϕ,θ, Ψ)one can define the overlap (or spatial Pearson correlation) *C*_*i*_ between the representation of unit *i* in A and that in B rotated by (ϕ,θ, Ψ). Assume that -1 < *C*_*i*_ < 1.

Then define *C*_*mean*_ (ϕ,θ, Ψ) as the mean *C*_*i*_ (ϕ,θ, Ψ) across all units, or the units in a cluster. Of course, -1 < *C*_*mean*_ (ϕ,θ, Ψ)< 1 as well (in practice its range is much more restricted, if different units do not coincide in their ‘best rotations’). Now position all Euler triplets along the x-axis given by their *C*_*mean*_ value, and define f(x) as their density (density of angles) along the axis. That is, if one considers a total of *N*_*angles*_, there are *N*_*angles*_ f(x)dx of them between x and x+dx. In Fig.S3A we show the *C*_*mean*_ distribution for an entire population (dashed line) and separately for each of 8 grid unit clusters (colored lines).

Assume now that among all *N*_*angles*_ angles, we pick for each unit *N*_*best*_ of them, those that have the highest *C*_*i*_ value. How will all the *N*_*best*_ × *N*_*units*_ angles be distributed, on average, in terms of f(x), the *C*_*mean*_ –ordered histogram? On average, they will concentrate more at higher *C*_*mean*_ values, at the very least because each unit gives a 1/N contribution to *C*_*mean*_ ; but possibly more concentrated than that. How much they concentrate is critical in order to determine the clustering coefficient *CC*, which measures simply how many of the *N*_*best*_ angles (what fraction) coincide among pairs of distinct units. We assume then that, at least within a cluster,

1. one can write:

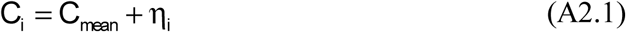

where η*i* is a form of ‘noise’, i.e. the combined effect of all other factors independent and unrelated to *C*_*mean*_. Note that this decomposition is a strong assumption.
2. this ‘noise’ is normally distributed, with a width σ(*N*_*best*_, cluster) that is the same across units in a cluster. If we denote with b(x; *N*_*best*_) the average fraction of *N*_*best*_ angles, among the *N*_*angles*_ f(x) present at a given *C*_*mean*_ –value x, such that, within a cluster,

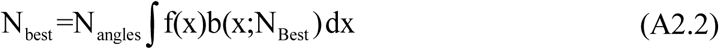

and with x_b_ (*N*_*best*_, cluster) the value of x such that this average fraction is ½, with these assumptions one has that b(x) can be expressed as the complementary error function

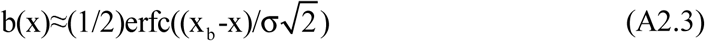

which Fig.S3B shows is not a bad approximation, if one allows x_b_ and hence b(x) to depend on both the cluster and *N*_*best*_. We make now the additional, critical (mean-field) assumption that
3. the clustering coefficient, *CC*, at least within each cluster, is only determined by the average density b(x; *N*_*best*_). Therefore, considering a generic pair of units,

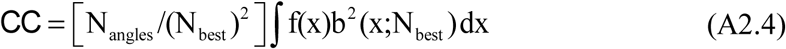 One may observe that CC is given by the extent of the overlap between the two distributions f(x) and b(x) (dropping for ease of notation its argument *N*_*best*_). Eq.(A2.2) and Fig.S3 show that for small *N*_*best*_ the overlap is limited to the opposing tails of the two distributions. We can evaluate the goodness of the assumptions made so far by comparing the values of the clustering coefficient obtained from the last equation to those obtained from the full analysis of simulation results. Equation A2.4 can be numerically evaluated by making use of the f(x) computed from simulations (Fig S3 A) and of the parameters for b(x) obtained from Gaussian fits (Fig S3 B-D). The resulting mean field approximation curve shows a remarkable similarity with simulation results (Fig 4 D). If we proceed and make the final assumption that
4. f(x) has a quasi-Gaussian upper tail

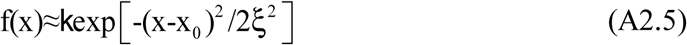

with k a suitable factor and ξ the effective width of the tail, we can obtain an analytical estimate of the *CC*. The result is

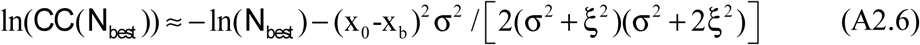

(keeping only the leading exponent). Both x_b_ and σ may depend on *N*_*best*_, but Fig.S3 indicates that within each cluster the dependence of σ is weak, while x_b_ shifts leftward as *N*_*best*_ increases: from Eq.(A2.2) one can derive

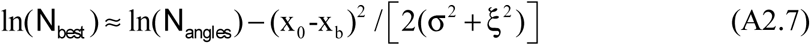 This yields a scaling of ln(*CC*(*N*_*best*_)) ≈ –β ln(*N*_*best*_) with 0 < β < 1, and the particular value β=1/2 is obtained for σ^2^ ≈ 2ξ^2^. We have no explanation for why this last relation appears to hold, approximately, for *N*_*best*_ small, as shown in Fig.4D.

## Supplementary Figures

**Figure S1.**
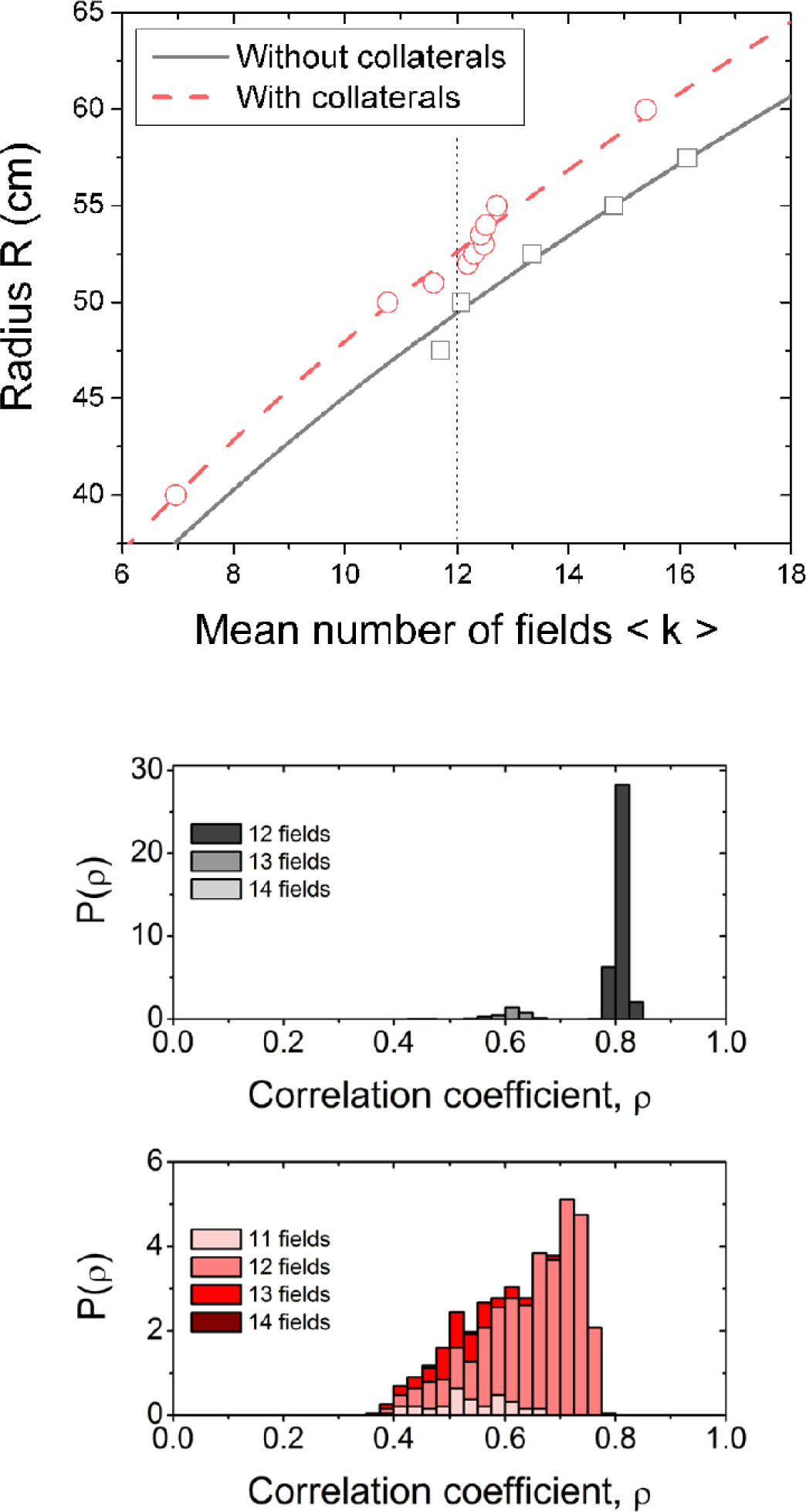
Interactions slightly reduce grid spacing, and distort the grids. Top. Dependence of the mean number of fields across units in each simulation on the radius R of the sphere, with (red) and without (grey) recurrent collaterals. Bottom. Distribution of the values of the spherical correlation of the map developed by each unit with the appropriately oriented perfect “soccer ball” pentagonal grid, in simulations with R=50cm (without collaterals) and R=52.6cm (with collaterals). Note that in the other “control” simulations without collaterals cited in the text, R was set to 52.6cm, and only the subset of units with 12 fields (a minority) was used in the analyses.

**Figure S2.**
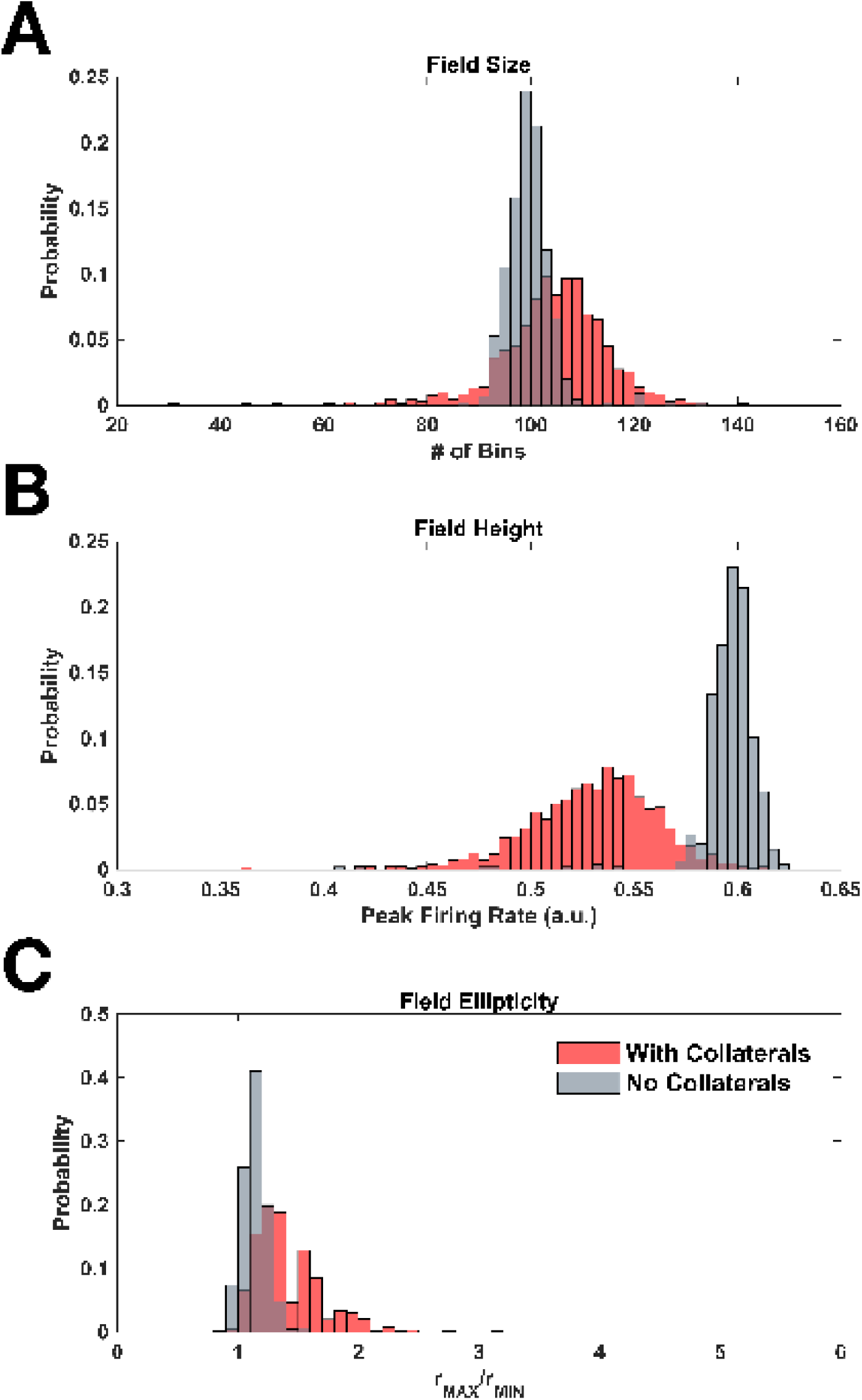
Interactions distort grid field shape. Distribution of field size (**A**), field maximum height (**B**) and field ellipticity (**C**) across a population of grid cells. Red bars: population with recurrent collaterals; grey bars: population without recurrent collaterals.

**Figure S3.**
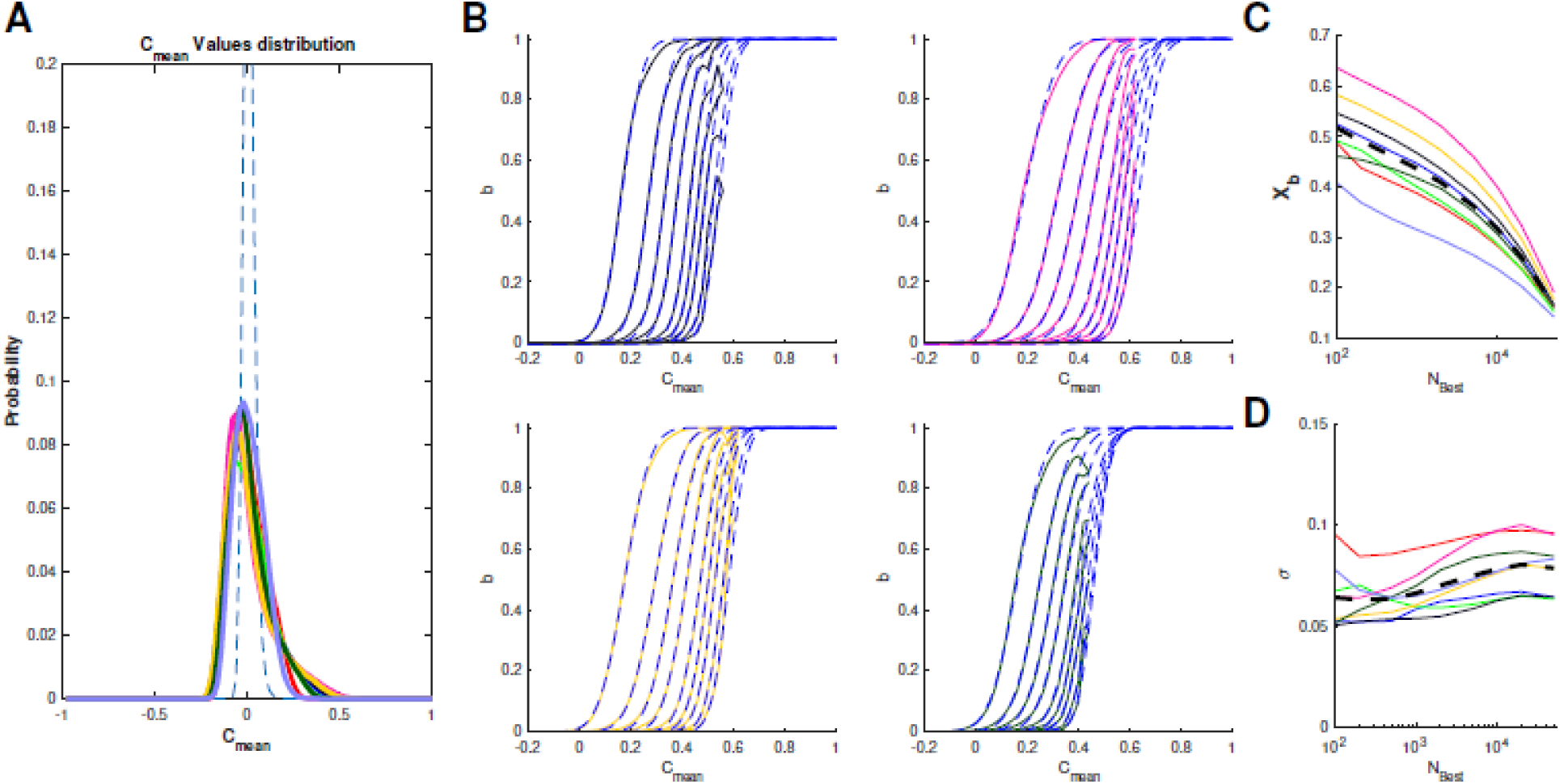
Despite the clustering, remapping appears to be a random process. A Density of C_mean_ values across the set of Euler rotations. Colored curves are computed within units belonging to the same cluster; the dashed line is the C_mean_ density computed over all units. B Four examples (4 different clusters) of the cumulative C_mean_ distribution computed for different values of N_Best_ [=100, 200, 500, 1000, 2000, 5000, 10000, 20000, 50000]. Colored lines: simulation data; dashed lines: Gaussian fits. C and D Mean and standard deviation, respectively, obtained from the Gaussian fits for different clusters and different N_Best_ values. Black dashed line: mean across clusters.

**Figure S4.**
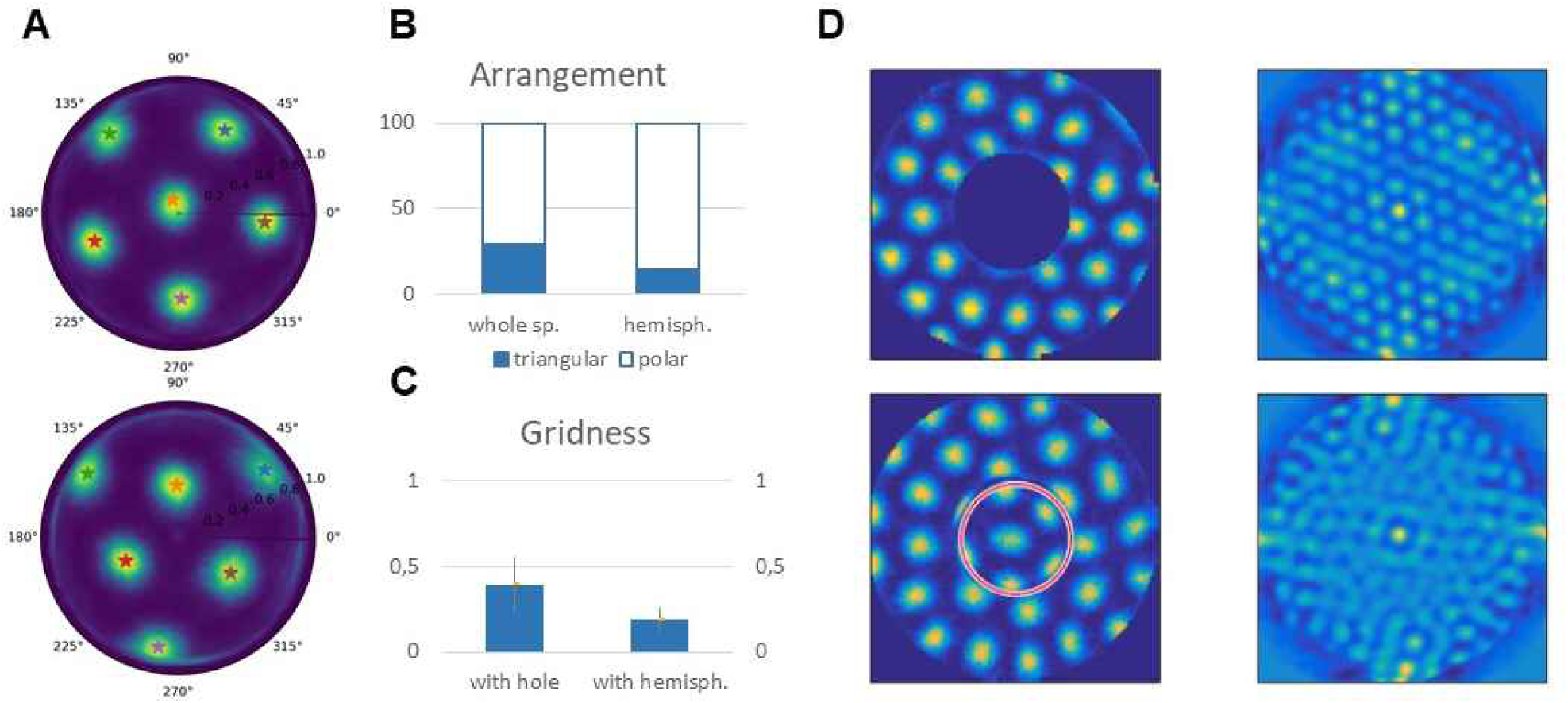
Boundaries repel grid fields and distort the grid also in curved environments. **A** Two possible arrangements of nearly regular grids on the hemisphere: the polar (top) and triangular (bottom). **B** Cutting in two halves a perfect soccer ball along an arbitrary equator, the triangular arrangement would occur with frequency 28.6%, whereas in simulations it occurs with frequency 14±5%. **C** Likewise, the average gridness score (computed on the flat surround alone) is reduced by the presence of a hemisphere in the center, relative to a hole. D Examples of grid maps emerging in the two cases, with their autocorrelations relative only to the flat surround.

**Figure S5.**
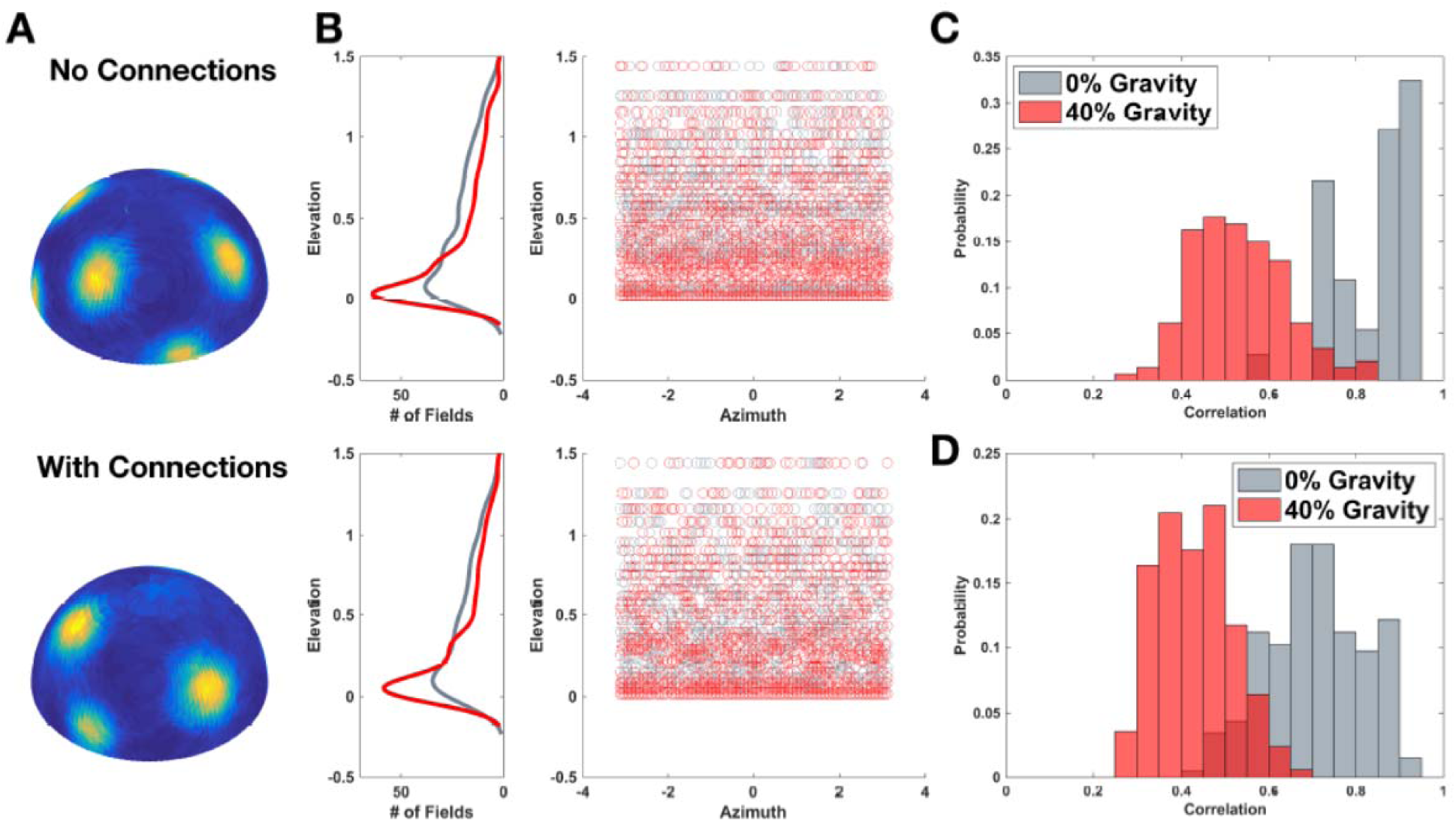
Gravity further distorts grids, over and beyond the effects of curvature. **A** Examples of grid maps emerging with ‘40% gravity’ (see text) without (top) and with (bottom) recurrent connections, as in the other panels of this figure. **B** Density of fields as a function of elevation (left; the apparent positive density below zero is due to smoothing) and position of each field from 100 units also along the azimuth (right). Red indicates the presence of gravity. **C** Gravity lowers the correlation with the perfect soccer ball, without interactions among units, from an average value of 0.84 to 0.53 (it would have been 0.93 on the full sphere). **D** With recurrent interactions, simulations show a similarly decreased correlation, to an average value of 0.43, from a lower baseline of 0.70 (higher than the value 0.66 of the full sphere), indicating approximately independent effects of the interactions and of gravity.

